# Identification of Gene Targets for the Sprouting Inhibitor CIPC

**DOI:** 10.1101/2024.12.07.626028

**Authors:** Thomas M Grand, James K Pitman, Alexander L Williams, Lisa M Smith, Andrew J Fleming

## Abstract

Sprout suppressants are widely used in industry to ensure year-round availability of potato tubers, significantly decreasing wastage by repressing premature growth of buds on the tuber surface during storage. Despite its ban from 2020 in the EU, isopropyl N-(3- chlorophenyl) carbamate (also known as chlorpropham or CIPC) remains the most widely used suppressant world-wide. However, the mechanism of action of CIPC remains obscure. Here we report on a combined targeted transcriptomic and genetic approach to identify components in the tuber bud cell-division machinery that might be involved in CIPC’s mode of action. This involved RNAseq analysis of dissected, staged tuber buds during in vitro sprouting with and without CIPC to identify lead genes, followed by the development and application of an Arabidopsis root assay to assess cell division response to CIPC in selected mutants. The ease of use of this model plant, coupled with its immense genetic resources, allowed us to test the functionality of lead genes encoding cell-division associated proteins in the modulation of plant growth response to CIPC. This approach led to the identification of a component of the augmin complex (a core player in mitosis) as a potential target for CIPC.

## Introduction

The potato tuber is a storage organ which has been bred to accumulate very large reserves of starch, leading to the crop being one of the most important sources of human nutrition world-wide and of significant agronomic and industrial value (Robertson et al. 2018). Ontogenically, the tuber is an enlarged stem, and on the surface of the tuber are a number of buds which are equivalent to apical and axillary meristems (Teper-Bamnolker et al. 2012). After maturing, the buds on the tuber are initially dormant (serving as an over-wintering device) but can subsequently be triggered to undergo growth, leading to sprouting as the meristem and sub-adjacent stem tissue rapidly grow and expand to establish a new plant. The ability to control sprouting over extended periods post-harvest is essential to the potato industry since premature sprouting leads to large-scale spoilage of the crop (Paul et al. 2016).

Most potato tubers have an innate eco-dormancy which delays sprouting for a variety-specific time, which can be extended via the control of environmental variables, e.g., light, temperature, and humidity (Sonnewald and Sonnewald 2014). However, the use of purely environmental factors to prolong tuber dormancy comes with associated challenges. For example, although lower temperatures help prevent sprouting, cold-induced sweetening can occur, leading to loss of commercial value through changes in taste, as well as the potential for enhanced acrylamide formation upon frying and associated potential health risks (Sowokinos 2001; Zhang et al. 2017). This trade-off between cold-induced sweetening and dormancy break can be alleviated by using chemicals to suppress sprouting, allowing higher storage temperatures. Indeed, without the use of chemical sprout suppressants, innate dormancy combined with environmental controls would not allow for the uninterrupted supply of potatoes between harvests to which the industry and end-users have become accustomed. However, some of the most effective sprout suppressants have come under increasingly stringent regulation in many countries, with isopropyl (3-chlorophenyl) carbamate (CIPC) (the most widely used sprouting suppressant world-wide) effectively banned in the EU since 2020 (European Food Safety (Authority 2017; Authority et al. 2017; Authority 2012). Although alternative suppressants are available, they are generally not as effective as CIPC and/or require extensive changes in storage infrastructure (AHDB (Board 2021). The development of new sprout suppressant chemicals and methods would greatly benefit the potato industry and end-users, limiting the large food wastage that will otherwise occur during tuber storage.

CIPC was first identified as a sprout suppressant 70 years ago (Marth and Schultz 1952), however our understanding of its mode of action remains very limited. Initial observations of abnormal mitotic spindles after CIPC treatment of roots led to the conclusion that CIPC prevents sprouting and plant growth by inhibiting mitosis (Eleftheriou and Bekiari 2000; Vaughn and Lehnen 1991). Further work revealed that it arrests cells at a prometaphase-like state, likely due to failure of spindle formation (Lloyd and Traas 1988) or at the G2/M checkpoint (Campbell et al. 2010). A general conclusion has been that CIPC acts as a mitotic disruptor, causing disruption of the microtubule organising centre (MTOC) and a loss of spindle polarity (Lloyd and Traas 1988; Vaughn and Lehnen 1991; Doonan et al. 1985; Traas et al. 1989). The mitotic spindle and phragmoplast appear to be particularly sensitive to CIPC, with the cortical arrays and interphase microtubules more resistant to disruption (Yemets et al. 2008; Bartels and Hilton 1973). In plants, CIPC is effective at doses in the µM order (Doonan et al. 1985) whereas any affects in mammalian cell requires treatment at concentrations of mM or higher (Holy 1998; Nakagawa et al. 2004). The lack of sensitivity observed in animal cells suggests that if CIPC is disrupting an element linked to mitosis, the target is, to some extent, specific to plant systems. However, the molecular identity of the plant-specific CIPC-target(s) remains unknown.

Previous efforts to understand its mode of action have been made using microarray profiling to identify changes in gene expression linked to CIPC-treatment of tubers. This approach identified changes in expression in a range of cell division-related genes, as well as those involved in response to oxidative stress and ABA (Campbell et al. 2010), however the functional significance of the lead genes identified in this study remains to be elucidated. Moreover, the microarrays available at the time represented only ∼10,000 potato genes while we know that the potato genome encodes approximately 39,000 proteins, thus some elements of the CIPC-related gene expression response were potentially missed. The advent of more powerful methods for analysing gene expression (RNAseq) from small amounts of tissue, coupled with much improved sequence resources, including anchor genomes, allows much deeper transcriptomic analysis, raising the possibility of analysing gene expression in individual tuber buds with or without CIPC treatment as sprouting progresses.

One of the challenges of research with potato is that the molecular genetic tools, resources and knowledge base are limited, particularly in comparison with the model plant *Arabidopsis thaliana*. However, it has become clear from investigations over the last two decades that much of the core molecular machinery involved in plant growth is highly conserved between angiosperms. This conservation particularly extends to the process of the cell cycle and division elements of cytokinesis (Han et al. 2022; Teper-Bamnolker et al. 2012; Campbell et al. 2010; Sablowski and Gutierrez 2022; Hartmann et al. 2011; Mouzo et al. 2022; Thoma and Zheljazkov 2022), raising the possibility of using lead information gained from one experimental system (potato) to inform experiments in another, more tractable system (Arabidopsis) to more rapidly test initial hypotheses, generating useful information to guide experiments in the original system. For example, it is clear that there is a directional cycle of kinase activity which drives plant cells through the classical elements of the G1, S, G2 and M phases, with specialised proteins (cyclins) playing a major role in controlling kinase activity, linking cell division to endogenous and exogenous signals. The most salient feature that distinguishes the plant cell division process from many other eukaryotes lies in the mechanics by which the two daughter cells are separated by a new cell wall. The site of this presumptive wall is marked by a unique sub-cellular structure (the preprophase band; PPB), with a new cell plate being formed internally, between the newly formed nuclei, which grows centripetally to join the mother cell wall at the sites previously marked by the PPB. This process involves the co-ordinated assembly and disassembly of cytoskeletal structures, and the agglomeration of vesicle structures containing cell wall material, both of which need to be timed with the classical formation and function of the mitotic spindle to allow correct separation of newly formed chromatids (Facette et al. 2019; Campbell et al. 2010). Although progress has been made in understanding the control of this process, the precise role of many components remains to be elucidated.

In this paper we report on a series of experiments designed to identify lead genes involved in the early response of potato buds to the sprouting inhibitor, CIPC. We describe the development of an *in vitro* sprouting assay which enabled us to collect and analyse dissected tuber buds at precise times after CIPC treatment, thus generating an analysis of gene expression using RNAseq over the first five days of CIPC response. This analysis is coupled to an Arabidopsis root growth assay which allowed us to test the functionality of genes identified in potato as putatively involved in the CIPC response. By comparing the CIPC-response of wild-type Arabidopsis roots with that of relevant mutants, we show that at least some lead genes from our potato bud analysis are functionally involved in modulating the cell-division response to CIPC.

## RESULTS

### CIPC suppresses growth of *in vitro* cultured buds

To establish an *in vitro* system to robustly test the effect of CIPC on tuber bud growth, we adapted the system reported by Hartmann and colleagues (2011). Buds were dissected from tubers, surface sterilised, then incubated on agar containing basal growth medium in multi-well plates with or without CIPC, as described in the methods section. Preliminary experiments showed that under control conditions (no CIPC) buds sprouted within about 5 days (sprout length at least 5 mm), whereas incubation with 100µM CIPC led to almost complete suppression of growth (sprouts less than 5 mm long even after 25 days incubation). To provide more detail on the differential growth response, we analysed the bud structure at different time points using SEM (**Fig. 1**). These results showed that at day 0 buds were sheathed by leaf primordia with the bud slightly sunken into the tuber surface (**Fig 1A**). By 24 hrs both the control (**Fig. 1B**) and CIPC-treated buds (**Fig. 1C**) showed some elongation of the covering leaf primordia but could not be reliably distinguished from each other. However, by day 5 there was a clear difference between control and CIPC-treated buds. New leaf primordia had formed in the control buds and there was clear elongation at the base of these leaves, lifting the tip of the bud clear of the adjacent tuber surface (**Fig. 1D**). In contrast, although new leaf primordia had also formed on the CIPC-treated buds, the primordia remained relatively compact with little elongation growth (**Fig. 1E**). This difference in extension growth was even more apparent by day 9, by which time the control buds had extended towards the top of the growth chamber, revealing an extensive elongated base on which hairs had formed (**Fig. 1F**). In the CIPC-treated buds at day 9, multiple leaf primordia were apparent, but they all remained stunted with little or no elongation growth away from the surface of the tuber (**Fig. 1G**).

**Figure 1.**
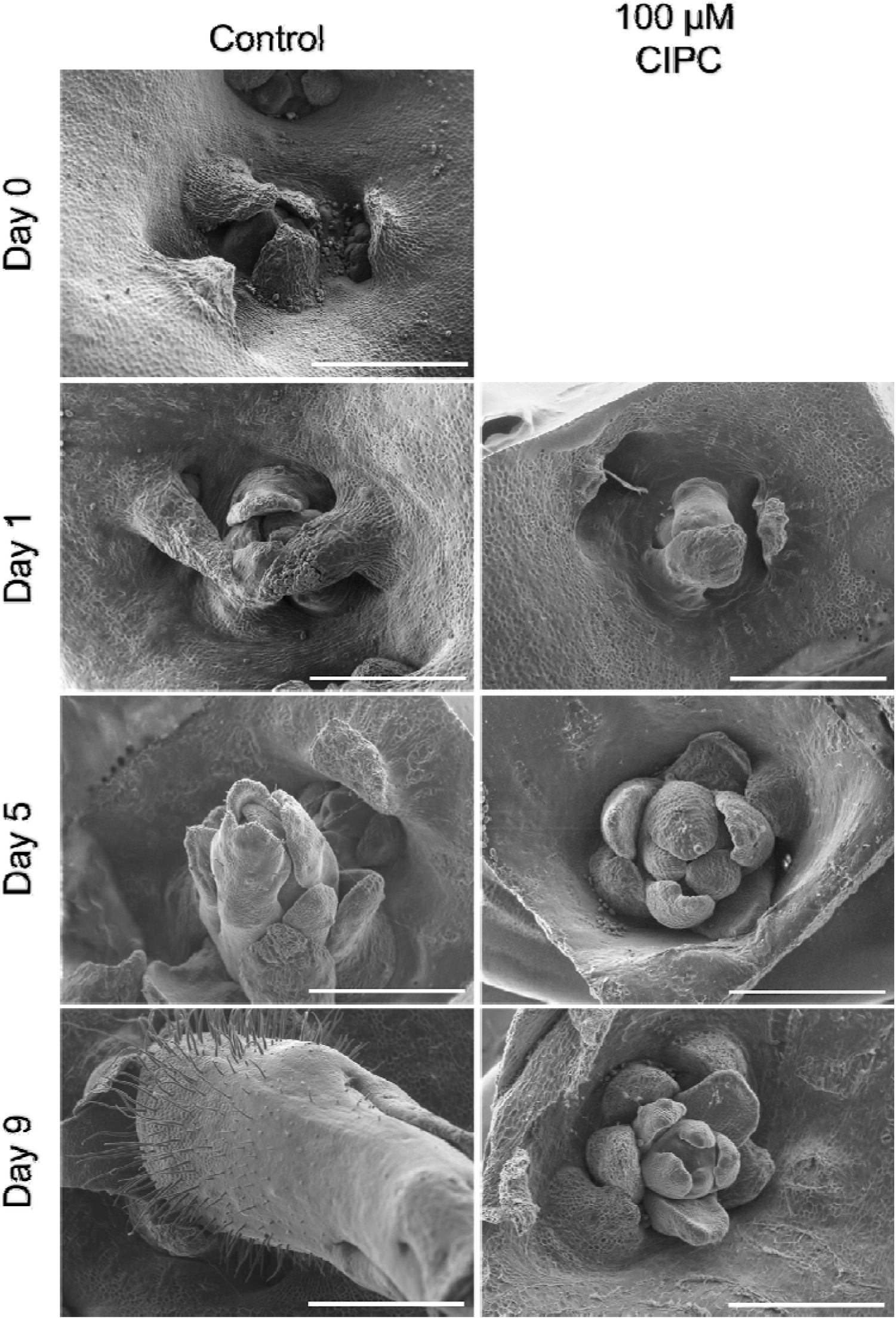
CIPC leads to early changes in bud morphology during sprouting SEM images of apical buds on the surface of a potato tuber at day 0 through to day 9 after sprouting initiation (as indicated) for explants grown without (A,B,D,F) or with (C,E,G) 50 µM CIPC. Scale bars = 1 mm.

### CIPC leads to changes in the transcriptome of buds by day 5 of treatment

To analyse the early impact of CIPC on gene expression, an RNAseq analysis was performed at different time points on pooled bud samples after growth with or without CIPC. As indicated in Fig. 1, the samples represented time points when there was no overt growth difference between treated and non-treated samples (day 1), a time point when there were clear growth differences at the micro-scale (day 5), and a time point when there were clear differences in growth at the macro-scale (day 9). Three biological replicates were taken for RNAseq analysis at each time point/treatment, with four dissected buds pooled for each replicate. Replicates of a day 0 untreated control sample were also included in the analysis. After initial cleaning of data and quality control, the RNAseq data were first analysed to check on the variation of expression pattern within and between treatments using a Pearson’s correlation coefficient matrix (**Fig S1**).

Biological replicates clustered closely with the one exception of replicate 3 of the day 0 control samples. This sample was distinct from the other day 0 control replicates but also did not group with the other samples. Due to natural variation in dormancy progression, it is possible the tubers in this sample were at a slightly different stage of dormancy. As this sample did not correlate with other sample groups, it was not excluded from subsequent analysis.

At day 1, both the control and CIPC-treated samples were highly correlated, however by day 5 the CIPC-treated replicates diverged in expression from the day 1 samples, while the control replicates did not. By day 9, CIPC-treated samples correlated most closely with day 5 CIPC-treated samples. From this initial analysis, it appeared that there was a transition point in gene expression response to CIPC between 1 and 5 days after treatment.

To verify the observations from the correlation analysis, principal component analysis (PCA) was used to investigate group similarity. The resulting plot (**Fig. 2**) confirmed that replicate samples typically grouped together well. By day 1 the CIPC and control samples had diverged from the day 0 controls but had similar expression profiles (compare green and red symbols). By day 5 the PCA enabled a clear discrimination between control and CIPC- treated samples (cyan symbols) and this discrimination was maintained in the day 9 samples (purple symbols). Interestingly, the CIPC-treated samples at day 5 and day 9 grouped together in the same vicinity of the plot as the day 0 controls. As with the data shown in Fig. S1, these results support the hypothesis that a major change in transcriptional profile occurs between day 1 and day 5 after CIPC treatment.

**Figure 2.**
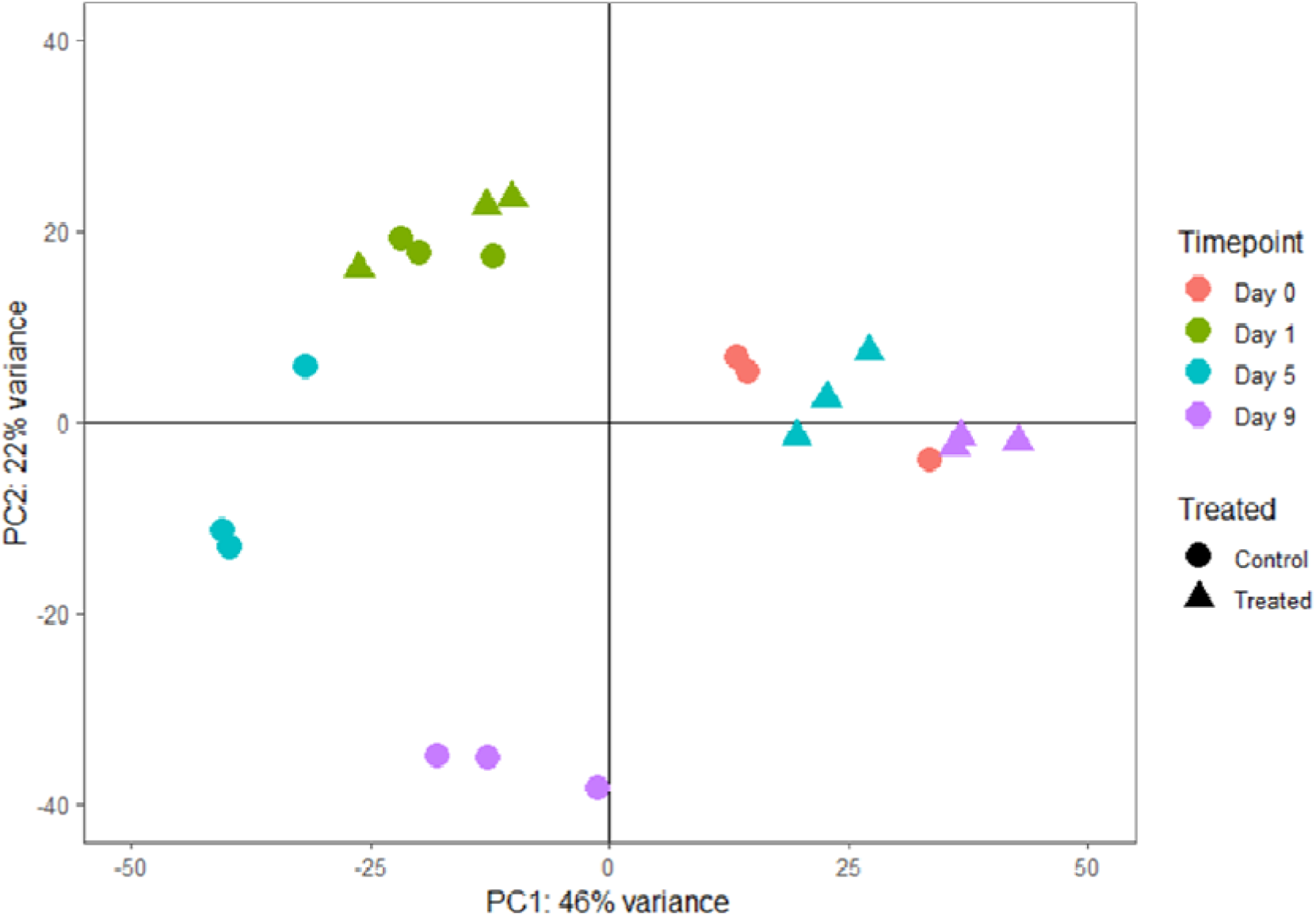
Principal component analysis of RNAseq data shows that CIPC-treated buds display distinct transcriptomes. PCA of RNAseq data obtained from either control (non-treated) (circles) or CIPC-treated (triangles) buds at time points (as indicated) after initiation of sprouting, with day 0 data from untreated buds. The first two principal components (PC1 & PC2) are plotted to show distinctions between sample groups, with each sample is plotted as a point. Samples are coloured according to time from sectioning and treatment. Clustering of point indicates a shift in the transcriptome of control samples with time that is distinct from the CIPC-treated samples.

### Genes related to cell division show decreased expression by day 5 after CIPC treatment

To explore the underlying changes in gene expression in more detail we performed K-means clustering of the 2000 genes showing the most variation in gene expression between treatments at each time point (**Fig. 3**). The data clustered into four groups, with the results visualised using a heatmap. The gene list for each cluster was analysed using gProfiler to identify over-represented GO terms.

**Figure 3.**
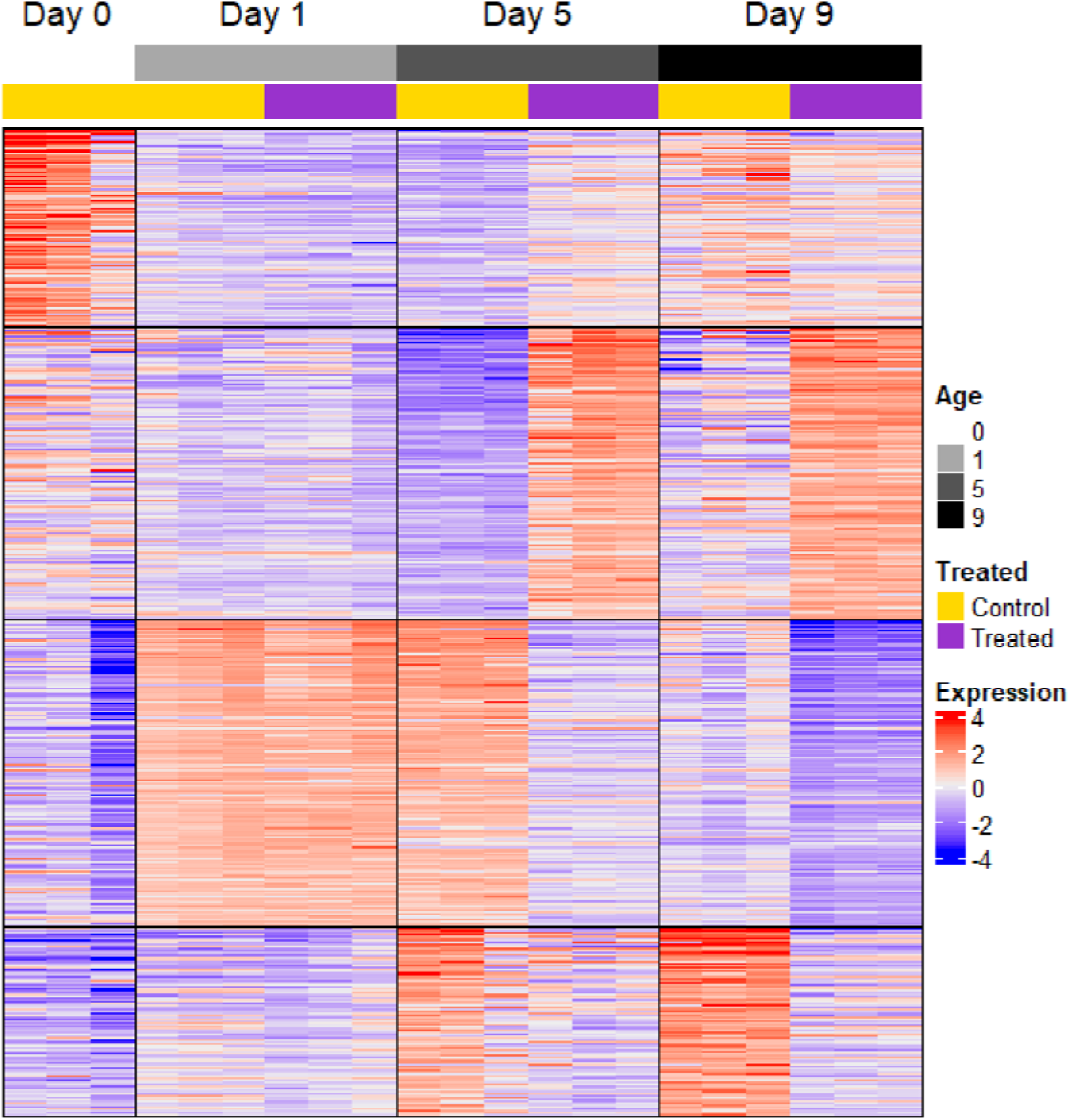
K-means clustering of RNAseq data indicates cell cycle-associated genes show a distinct pattern of expression in CIPC-treated buds compared with controls within 5 days of sprouting initiation. The top 2000 genes from RNAseq analysis showing the most variable expression (analysed using SESeq2) were grouped using K-means clustering to detect patterns of expression. Data have been scaled and plotted as a heatmap, as indicated by the scale. Datasets are defined by age (days after initiation of sprouting) and whether they represent untreated control or CIPC-treated buds (as indicated). Four clusters were identified: A - Abiotic stress response; B - Toxic substance response/hydrogen peroxide catabolism; C - Cell cycle/mitosis; D -Photosynthesis. Expression was calculated by normalising gene counts with the Variance Stabilising Transform, centring by subtracting mean expression for each gene, and scaling by dividing by gene standard deviation.

Cluster A consists of 404 genes and is enriched for GO terms relating to regulation of abiotic stress response (P_adj_ = 3.47×10^-12^) and biosynthetic processes (P_adj_ = 3.48×10^-2^). This cluster was highly expressed at day 0, with minimal differences between control and CIPC-treated samples at later time points. Within these high-level GO terms, the biosynthetic processes GO term includes genes related to RNA biosynthesis and regulation of transcription, while the stress response GO term includes genes involved in osmotic stress, cold stress response and response to hydrogen peroxide. This cluster therefore represents genes that were active during storage or sampling, which were then downregulated during the experimental time course.

Cluster B includes 589 genes and is enriched primarily for oxidation-reduction processes (P_adj_ = 2.721x10^-12^) and hydrogen peroxide catabolism (P_adj_ = 5.612×10^-7^) as part of the cellular detoxification GO term family. These genes were generally upregulated in the CIPC- treated samples and downregulated in the control samples. The gene list includes a number of peroxidases and catalases which are likely expressed in response to an increase of reactive oxygen species in response to CIPC.

Cluster C contains 625 genes and is enriched for cell cycle-related genes (P_adj_ = 3.237×10^-28^), microtubule-based processes (P_adj_ = 2.762×10^-24^) and steroid biosynthetic processes (P_adj_ = 1.122×10^-5^). These genes were upregulated at day 1, with expression diverging between the control and CIPC treated samples by day 5. The expression of these genes (encoding proteins involved in many aspects of cell division, such as cyclins, kinesins, tubulins, transcription factors and cell wall biosynthetic enzymes) was downregulated in the CIPC-treated samples but remained upregulated in control samples. This aligns with decreased cell division in CIPC-treated buds compared to rapidly growing control samples.

Finally, cluster D consists of 382 genes and is enriched for photosynthesis related genes (P_adj_ = 7.110×10^-74^) and negative regulation of endopeptidase activity genes (P_adj_ = 3.26x10^-14^). Genes in cluster D were upregulated in expression in the control samples from day 1 onwards, while expression in CIPC-treated samples remained stable. These results fit to the observed development of green leaf primordia in the control samples whereas in the CIPC- treated buds the emerging leaves appeared abnormal and retarded.

Taken together, the data in Fig. 3 were consistent with the hypothesis that changes in the expression of genes involved in cell division (cluster C) played a major role in distinguishing the early response to CIPC by day 5 after treatment.

In parallel to the analysis of the 2000 genes showing the most variable expression across all samples shown in Fig. 2, we performed a broader analysis of gene expression response to CIPC in which DESeq2 was used to identify all differentially expressed genes (DEGs), defined by those showing at least a two-fold change of expression and an P_adj_ < 0.05. In total, 12,437 DEGs were identified by comparison of different timepoints for each treatment, of which 2,338 were unique to the control samples, 2,639 were unique to the CIPC-treated samples and 7,460 were differentially expressed in both time courses. GO terms associated with DEGs were then identified by using the Plant GO Slim subset of GO terms with annotations from AgriGO v2 (Tian et al., 2017) and GOslimmer (Faria, 2019), followed by enrichment analysis through g:Profiler.

After removal of redundant terms, 26 GO terms were identified with differential expression. In the earliest phase of sprouting (from day 0 to day 1), there was wide downregulation of cell communication-related processes alongside upregulation of metabolic and growth-related processes in the control samples. Photosynthesis, generation of precursor metabolites and energy, and carbohydrate metabolic processes were upregulated early in the time course and remained upregulated, fitting with a narrative of storage mobilisation and development of photosynthetic competency early in bud growth. Gene expression levels were relatively stable in control samples between day 1 and day 5, with the exception of some stress/stimulus response processes, potentially signalling the end of wounding response and acclimatisation to new growth conditions. From day 5 to day 9 there was a partial reversion of the trends seen early in sprouting, with upregulation of cell communication and downregulation of cell cycle-related genes.

In comparison to the control samples, there were three major differences in gene expression identified in response to CIPC; earlier upregulation of cell communication genes, earlier downregulation of cell cycle genes, and a peak in expression of translation-related genes at day 1. Changes in expression of cell cycle- related genes were among the strongest changes elicited by CIPC from this analysis, with CIPC treatment leading to a strong downregulation of cell cycle-related genes between days 1 to 5. Since our RNAseq data supported the hypothesis that plant cell division is a target for CIPC, we focussed on a deeper analysis of this gene set.

Gene sets corresponding to the cell cycle GO term were compared between the control and CIPC time courses to assess overlap of the upregulated genes from day 0 to day 1 and downregulated genes from day 1 onwards. From day 0 to day 1, 134 DEGs (∼88%) related to the cell cycle were shared between the two treatments, with sixteen DEGs unique to the CIPC-treated samples and two unique to the control samples (**Fig. 4A**). Broadly, the same set of DEGs were downregulated later in the time course; 111 cell cycle-related genes were downregulated in the control samples from day 5 to day 9, with 19 unique to the control samples (**Fig. 4B**) and 92 shared with the CIPC-treated samples. Interestingly, 45 genes were uniquely downregulated in the CIPC-treated samples, with the majority (39 genes) downregulated by day 5 (**Fig 4**). These results are consistent with the hypothesis that CIPC treatment leads to an earlier downregulation in expression of cell cycle genes than occurs in control buds, and that it leads to altered expression of a broad yet specific set of cell cycle-related genes. In particular, the 45 genes that are uniquely downregulated in CIPC-treated samples can be viewed as lead candidates encoding proteins which may be closely involved in the early repression of cell division observed after treatment with CIPC.

**Figure 4.**
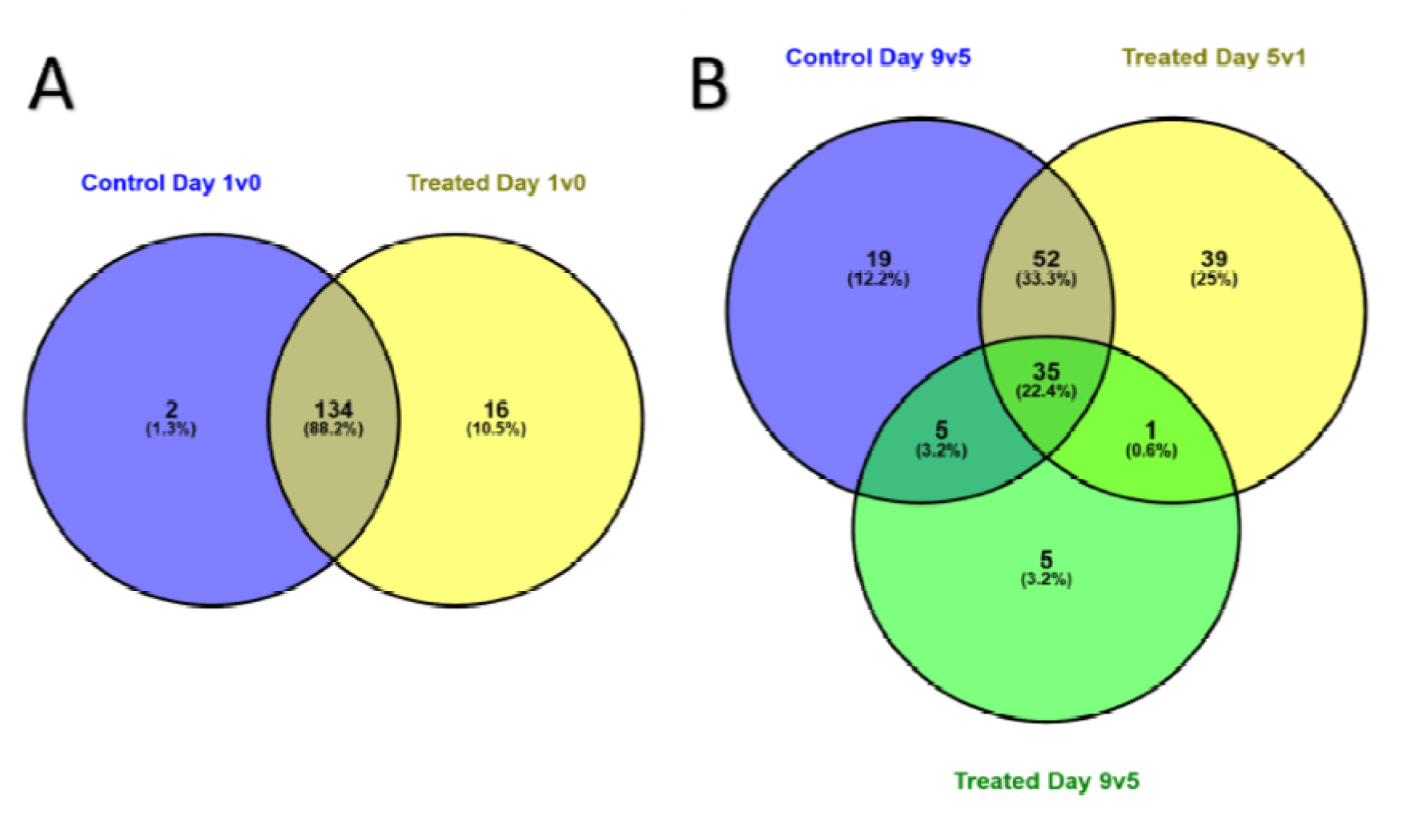
Comparison of cell cycle associated genes identifies 39 whose expression is down-regulated within 5 days of CIPC-treatment compared with control. Comparison of gene sets corresponding to GO-Cell Cycle term in RNAseq data from control and CIPC-treated buds at 0, 1, 5 and 9 days after sprouting initiation, as indicated. Values give absolute number of genes, with percentage indicating the relative contribution within the comparison made.

### A CIPC-response assay in Arabidopsis allows functional testing of lead cell cycle-related genes linked to bud suppression in potato

To enable a relatively rapid initial functional test for lead genes linked to the repression of the cell cycle identified via RNAseq analysis of CIPC-treated potato buds, we developed a CIPC-growth response assay using Arabidopsis roots. WT (Col-0) seedlings were grown vertically in a standard growth medium and root length measured over time. By day 7, roots had grown to approximately 26 mm (**Fig. 5A**). When CIPC was included in the medium, growth was inhibited at all concentrations above 1 µM, with growth being progressively more repressed with higher CIPC concentration (**Fig. 5A**). Since inclusion of 10 µM CIPC in the medium led to a consistent repression of root growth but nevertheless allowed some measurable growth to occur, we performed a screen of T-DNA mutants in Arabidopsis using 10 µM CIPC. We also included a 20 µM CIPC-treatment in the screen to ensure a strong growth repression in our analysis.

**Figure 5.**
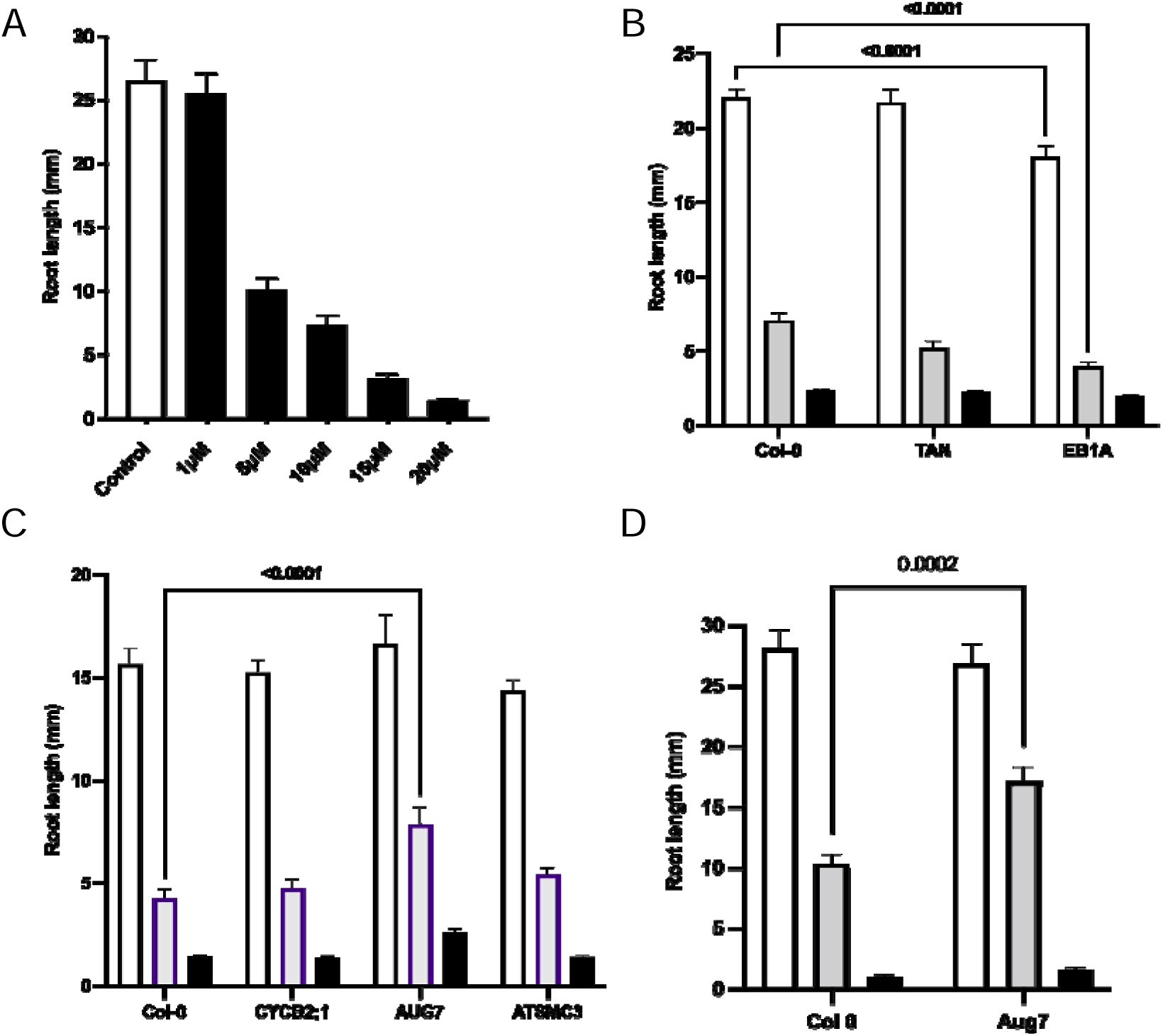
An Arabidopsis root growth assay identifies lead cell cycle genes functionally involved in the CIPC-response **(A)** Root lengths of Col-0 seedlings at 7 days after germination on medium containing a range of CIPC concentration, as indicated. **(B-D)** Root lengths of T-DNA mutants for genes identified as potentially involved in CIPC growth suppression. For each set of T-DNA mutants analysed, Col-0 seedlings were assayed in parallel. Seedlings were germinated on either 10µM (grey bars) or 20 µM (black bars) CIPC and root length measured after 7 days. Two-way ANOVA was carried out with Dunnet’s multiple comparison test for each assay, with samples showing a difference in growth after 7 days at a particular CIPC concentration indicated where p < 0.001. Bars show mean with standard error of the mean. For A-D, n > 34 per sample per treatment.

To select Arabidopsis mutants, we identified orthologues of the potato cell cycle genes identified by our expression analysis (**Fig. 4**), obtained relevant mutants from stock centres, confirmed the mutation by genomic PCR analysis, then performed root growth analysis. To allow for observed plate-plate variation in absolute root growth, we included Col-0 seedlings in each plate, with these values being used as the control within each dataset collected.

A total of 23 cell-cycle-related mutants were confirmed by genotyping and analysed (**Table S1**). Although some showed a potential altered growth response to CIPC, an altered growth of the root in the absence of CIPC complicated assessment. For example, as shown in **Fig. 5B**, loss of EB1A (a regulator of microtubules) led to a general decrease in root growth, making it difficult to assess any CIPC-specific element of further growth repression. Consequently, of the 23 mutants analysed, only one showed a strong and reproducible altered growth in the presence of CIPC with no evidence of altered basal root growth. AUGMIN7 encodes a component of the Augmin complex which has been shown in other eukaryotes to be required for correct mitotic spindle formation (Romeiro Motta et al. 2024; Hotta et al. 2012). The *aug7* mutant displayed significant loss of growth repression in the presence of 10 µM CIPC (P < 0.0001) (**Fig. 5C**), a phenotype that was reproducible in independent experiments (P = 0.002) (**Fig. 5D**). To explore whether mutations in any other genes (non-cell cycle related) displayed similar CIPC-related growth phenotype (to gain an estimate of the background frequency of altered CIPC-response in Arabidopsis mutants) we analysed a further 31 confirmed T-DNA mutants, selected on the analysis of DEGs between control and CIPC-treated samples, but which did not encode cell cycle-related proteins (**Table S2**). None of these mutants displayed a significant difference in CIPC response without showing a significant difference in growth on media without CIPC.

## DISCUSSION

Research over many decades has suggested that CIPC exerts its influence as a sprout suppressant via disruption of the cell division process (Eleftheriou and Bekiari 2000; Vaughn and Lehnen 1991). The results reported in this paper support this hypothesis and extend previous work by identifying a series of cell cycle-related genes whose expression is altered in the earliest phase of the CIPC growth repression response. Moreover, by exploiting the evolutionary conservation of cell cycle machinery, we could use an Arabidopsis root growth assay to provide functional data indicating that at least one of the lead genes identified plays a role in modulating the cell division response to CIPC.

### An *in vitro* system for potato bud growth allows analysis of the earliest events of the CIPC response

Our system for analysing tuber bud growth *in vitro* was adapted from that established by Hartmann and colleagues (2011). Coupled with the use of freshly grown tubers, it allowed for a highly standardised and reproducible process of bud sprouting. The *in vitro* system involves damage to the tissue around the explanted bud which does not occur during sprouting in intact tubers. Indeed, our analysis identified a swathe of gene expression responses linked to abiotic stress during the first 24 hrs after excision and incubation (**Fig. 3**). It is difficult to distinguish the extent to which this aspect of the transcriptional response is an artefact of the experimental system or gives a true indication of an element of the normal sprouting response. Importantly (as discussed below) other portions of the transcriptional response encompassed elements that clearly aligned with the observed and expected growth response of bud sprouting (cell division, photosynthesis). As long as the changes potentially linked to the *in vitro* element of the explanted bud growth are taken into account, we believe that the system described here provides a robust and rapid system for analysing the early phase of tuber bud sprouting.

With respect to CIPC, the *in vitro* system allowed observations on the early morphological response of buds treated with this sprouting inhibitor. Although CIPC-treated buds did generate primordia, their growth was greatly retarded and abnormal compared with control, untreated buds (**Fig. 1**). Our data indicate that CIPC elicits growth repression within 5 days of treatment under the conditions described here. Importantly for this investigation, the robust and reproducible bud sprouting (with and without CIPC) allowed the collection of samples for RNAseq analysis from buds that were at distinct developmental stages, giving us confidence that we were capturing gene expression responses to CIPC treatment of relevance to the growth response observed. The reproducibility of the differences in overall gene expression observed at the three time points analysed with the two treatments (control vs CIPC) (**Fig. S1**), provide confidence in the utility of the *in vitro* growth assay, in the technical quality of the RNAseq data and, consequently, in the interpretation of these data, described below.

### RNAseq analysis allows identification of lead genes involved in CIPC-induced growth repression in buds

Early investigations aimed at characterising the gene expression response of tubers used microarrays (Campbell et al, 2010). Although powerful, these approaches lacked the whole genome approach facilitated now by the development of newer techniques to analyse gene expression (RNAseq), and the larger and better annotated genome sequence databases against which such expression data can be anchored. Liu et al (2015) performed an RNAseq analysis of dormancy break in tubers using RNA extracted from parenchyma, with their 70% read mapping rate indicating the feasibility of RNAseq in potatoes. The overall consistency of results between our triplicate samples of each time point and treatment indicates robustness of the assay despite the potential for differences in dormancy stage between tubers.

The results reported here provide an advance on earlier microarray studies by providing gene expression data at a global level during the earliest stages (days) of the CIPC response. At a broad level, they indicate that during the first 24 hours of treatment it is difficult to distinguish buds based on gene expression pattern, but that by 5 days there are clear major differences between CIPC treated and control samples (**Fig. 2**). This correlates with the timing of the first detectable changes in growth response revealed by SEM analysis (**Fig. 1**). With respect to the type of genes showing major changes in gene expression at this stage, those annotated as cell cycle-related stood out as being repressed relative to the control buds, fitting to the hypothesis that CIPC is targeting the plant cell cycle machinery to repress growth. There was also a difference in the expression of genes related to photosynthesis, which we interpret as being a consequence of CIPC-related growth retardation leading to disruption of photosynthetic differentiation. During normal sprouting visible greening of the sprout occurs as it extends towards the light. This did not occur in the CIPC-treated buds.

The other major families of genes showing altered expression in our system related to stress response and toxin metabolism. As indicated above, we suspect a portion of these reflect the damage involved in explanting the buds, with the increase in toxin metabolism related to the fact that a xenobiotic (CIPC) was added to the growth medium. The identification of these gene responses and their interpretation as an intrinsic by-product of the experimental protocol means that they can, to a large extent, be discounted and effort focussed on those gene expression changes more likely linked to the biological processes occurring during bud sprouting.

With respect to cell cycle-related genes showing a differential expression response to CIPC, our analysis led to the identification of 46 genes, with the majority (40) being linked to events between 1 and 5 days after CIPC treatment (**Fig 4**). The plant cell cycle is highly homologous to that observed in other eukaryotes, with many hundreds of proteins implicated in the mechanics and regulation, depending on species (Sablowski and Gutierrez 2022; Polyn et al. 2015). The cell cycle-related genes identified here thus clearly represent a subset of the total number of genes involved in the plant cell cycle. Consideration of the types of cell cycle genes identified reveals a broad swathe, ranging from various cyclins involved in regulating transition around the cycle, through proteins involved in chromatin structure, to cytoskeletal elements implicated in cytokinesis.

If CIPC does function by impinging on the cell division process, then altered expression of many cell cycle-related gene can be envisaged as part of the system response to a major disruption of an essential process. Although a number of the genes identified have been demonstrated to functionally alter plant cell division and growth (Sablowski and Gutierrez 2022), it is difficult to assign a direct function in the CIPC-response (as opposed to altered expression being part of an indirect response to CIPC). To address this challenge, we developed a root growth assay in Arabidopsis to measure CIPC response.

### Use of Arabidopsis root growth to test CIPC response

The eukaryotic cell cycle is highly conserved. We therefore reasoned that if CIPC is interacting with a cell cycle-related component in potato buds, it is likely that a similar cell cycle component will be present in other plants. Moreover, due to the fundamental linkage of cell division to growth in most circumstances, CIPC would lead to altered cell division and growth in organs other than buds in other plants. To explore this possibility, we adapted an Arabidopsis root growth assay, a system which has been successfully used to explore the control of root growth (including the role of cell division) (MulJssig et al. 2003). Our results demonstrate that Arabidopsis roots show clear CIPC growth repression, with growth almost totally absent when seeds are grown in the presence of 20 µM CIPC. At lower concentrations (10 µM CIPC) some root growth does occur, but far less than observed in control, untreated seedlings. Arabidopsis does show CIPC-mediated growth suppression, thus it seems plausible that the targets for CIPC growth repression in Arabidopsis and potato are similar.

By identifying orthologues in Arabidopsis of the cell cycle-related genes identified in potato buds in our analysis of CIPC response, we were able to exploit the large and publicly available genetic resources to obtain a range of T-DNA mutants in the respective genes. A number of these genes had already been relatively well characterised and implicated in the normal process of plant growth and division, e.g., CYCD3 (Dewitte et al. 2007); TAN and RanGAP1 (Mir et al. 2018; Müller et al. 2006); CYCA1:1 (Fülöp et al. 2005). However, it should be borne in mind that even in Arabidopsis, genetic redundancy of cell cycle-related genes means that phenotypes in single mutants are often very limited (Dewitte et al. 2007). Bearing this out, the majority of the cell-cycle mutants analysed here did not show a significant shift in root growth in the absence of CIPC. When CIPC was included in the medium, the vast majority of the cell-cycle mutants did not show a significant change in root growth when a stringent confidence limit was set. The one exception to this was a mutant in a gene encoding AUGMIN7 (*aug7-2*), a component of the augmin complex which is involved in microtubule initiation in the spindle and phragmoplast in plants (Hotta et al. 2012) and for which there is extensive data in other eukaryotes for a major role in spindle function during mitosis (Lee and Liu 2019). The augmin complex has microtubule binding activity and localises gamma-tubulin to existing microtubules to initiate microtubule-dependent microtubule nucleation (Song et al. 2018; Hotta et al. 2012; Lee and Liu 2019). Gamma-tubulin no longer localises to the spindle or phragmoplast in *aug7* knockdowns and the spindles are long and disoriented (Hotta et al. 2012).

Our data implicate AUGMIN7 in CIPC-mediated growth suppression. Further work is needed to investigate whether this is an indirect role or whether, potentially, AUGMIN7 itself is a direct target. Augmin is a complex comprised of eight subunits, of which AUGMIN7 is one. It is intriguing to speculate that if AUGMIN7 were a target for CIPC, it might lead to a functionally compromised complex in which the resultant growth suppression would only be observed if AUGMIN7 were present, thus accounting for the decrease in CIPC-mediated growth suppression in the *aug7-2* mutant. In such a scenario, the lack of growth phenotype in the *aug7-2* mutant in the absence of CIPC would require a level of functional redundancy with the other subunits of the AUGMIN complex.

In summary, our results describe both a platform for the identification of genes involved in the early response of tuber buds to the sprouting inhibitor CIPC, and provide an analysis of lead cell cycle genes that modulate plant response to CIPC. Deeper understanding of the mode of action of CIPC, the most effective known suppressor of bud growth, may identify related, novel targets for the suppression of bud sprouting.

## Acknowledgements

Immense thanks go to Glyn Harper, Adrian Briddon (from AHDB’s Sutton Bridge Crop Storage Research (closed December 2021)) and Alice Sin (AHDB) for their support and advice throughout the project, as well as the provision of initial plant material (tubers) and chemicals (CIPC).

## Author contributions

TMG, JKP, AJF and LMS planned the experiments; TMG performed most of the experiments and analysed the data; ALW performed the remaining experiments and data analysis; AJF and LMS supervised the project; AJF, LMS and TMG wrote and edited the manuscript.

## MATERIALS AND METHODS

### Plant lines and growth conditions

*Solanum tuberosum* (King Edward cultivar; purchased from Suttons Consumer Products) were propagated in glasshouses in 15 L pots in Levington M3 compost. Plants were grown with a photoperiod of 12 hours, 20°C day/12°C night. Supplementation with 200 µM fluorescent light occurred if ambient light fell below 1000 µM. Tubers were harvested upon plant senescence, and stored in paper bags (dark, 6°C) for 12 weeks.

### In vitro sprouting assays

12-well plates (Cellstar) were prepared with 2 mL autoclaved media per well (½ Murashige and Skoog Basal Media (SIGMA), 1% w/v plant agar (Duchefa Biochemie), pH 5.8). 1 mM CIPC (SIGMA) was dissolved in methanol and added to a final concentration as specified.

Tubers were washed in tap water, surface sterilised in 1% NaClO for 10 mins, then rinsed with running tap water. Cores surrounding the apical were excised with a #12 cork borer (18.75 mm diameter) and trimmed to 5 mm depth. The excised buds were washed three times in distilled water, then transferred to the 12-well plate. Plates were then kept at 22°C in the dark.

### Scanning Electron Microscopy

Dissected buds were mounted onto aluminium stubs using OCT (Optimal Cutting Temperature compound, Sigma Aldrich), then imaged using a TM3030 Hitachi Tabletop Scanning Electron Microscope. Images were taken in secondary electron mode at 15kV. Cell size measurement was carried out in ImageJ (Schneider, Rasband and Eliceiri, 2012).

#### RNAseq transcriptomes and analysis

Potato buds were sprouted using the assay described above, on control and 100 µM CIPC agar with samples taken at days 0, 1, 5, and 9. Tissue was pooled from four buds for each biological replicate. RNA was extracted using the Sigma Plant Total RNA kit (Sigma Aldrich). RNA quality was assessed using gel electrophoresis, a Nanodrop 2000 (Thermo Scientific), and a Qubit broad range RNA assay kit (Invitrogen), as per manufacturer’s instructions. RNAseq library preparation and sequencing was performed by Novogene (https://en.novogene.com/). Samples were enriched for mRNA using polyA capture, and library preparation was with the NEB Next® Ultra™ RNA Library Prep Kit. Sequencing was performed on an NovaSeq 6000 S4 to a depth of at least 25 million paired end reads per library, read length 150 bp. All samples passed FastQC quality controls, with most reads having a Phred score of 36 or greater. Raw reads were mapped to an index built using

HISAT2 (Kim et al. 2019), based on the *S. tuberosum* doubled monoploid assembly version 4.03 pseudomolecule sequence and corresponding gene annotation (Hardigan et al. 2016). Mapped reads were mapped to each genomic feature using featureCounts (Liao et al. 2014). Mapping rates were consistent across samples.

RNAseq count data was filtered for genes with active expression (at least 3 samples with 10 counts each). Expression of these 27302 active genes was normalised in DESeq2 using the variance stabilising transform (VST) function. To explore the high-level similarities in intra- and inter-sample group comparisons, a correlation matrix of sample distances was created by calculating sample distances from the VST normalised dataset, with removal of genes with low expression variability (bottom quartile) to exclude stably expressed genes. Pearson’s rank correlation coefficients were calculated for each sample combination. Sample combinations were assigned a value from -1 to 1 from perfect negative correlation to perfect positive correlation. To further investigate sample grouping, principal component analysis (PCA) was performed on the top 500 most variable genes of the normalised dataset. A biplot of the first two principal components was created using ggplot2.

To detect patterns of expression, a heatmap of the top 2000 variable genes was generated. Values were centred by subtracting mean expression for each gene and standardised by dividing by gene standard deviation. Genes were grouped into clusters using K-means clustering. The number of clusters was determined by identifying the inflection point in an elbow plot. Clustered genes were analysed on g:Profiler to detect overrepresented gene ontology (GO) terms (Raudvere et al. 2019). Functional analysis by g:Profiler uses a GO term specific algorithm to set significance thresholds and to calculate an adjusted P value, correcting for multiple testing.

For differential gene expression analysis, the RNAseq dataset was analysed using DESeq2 (Love et al. 2014), modelled using a modified matrix from design = ∼Age*Treated, with day 0 as the reference level. Comparisons were called using the lfcShrink() function. The results were filtered for genes with greater than two-fold change and an adjusted p-value of less than or equal to 0.05. Results were ordered by adjusted p-value and separated into up- and down-regulated genes. Gene overlap was visualised using Venny (Oliveros 2007). The ordered gene lists were used for functional enrichment analysis on gProfiler (Raudvere et al. 2019) by converting to GO terms and identifying overrepresentation. The GO terms and associated p-values from functional analysis through gProfiler were processed using Revigo (Supek et al. 2011) with an allowed similarity level <0.5. Revigo reduces GO term list length by removing redundant and dispensable terms.

