## Supplemental table 1 for "Identification of Gene Targets for the Sprouting Inhibitor CIPC"

**Table S1**: Arabidopsis cell-cycle related mutants tested for altered root growth response to CIPC

| **Orthologue name** | **T-DNA line ID** | **Locus** | **Description** |
| --- | --- | --- | --- |
| ATSMC3 | SALK_204716C | At5g48600 | Structural maintenance of chromosomes protein |
| AUG7 | SALK_092584C | At5g17620 | AUGMIN subunit 7 |
| CDC6B | SALK_128156C | At1g07270 | Cell division control protein 6 homolog B |
| CMT3 | SALK_148381C | At1g69770 | DNA (cytosine-5)-methyltransferase CMT3 |
| CYCA1-1 | SALK_045486C | At1g44110 | Cyclin A1-1 |
| CYCB2-1 | SALK_019772C | At2g17620 | Cyclin B2-1 |
| CYCB2;4 | SALK_073438C | At1g76310 | Cyclin B2-4 |
| CYCB2;5 | SALK_207168C | At1g20590 | Cyclin B2-5 |
| CYCD2;1 | SALK_049449C | At2g22490 | Cyclin D2-1 |
| CYCD3;1 | SALK_045277C | At4g34160 | Cyclin D3-1 |
| CYCD3;2 | SALK_031739C | At5g67260 | Cyclin D3-2 |
| CYCD3;3 | SALK_207843C | At3g50070 | Cyclin D3-3 |
| CYCD4;2 | SALK_127016C | At5g10440 | Cyclin D4-2 |
| EB1A | SALK_138692C | At3g47690 | Microtubule-associated protein RP/EB family member 1A |
| FTSZ1 | SALK_046941C | At5g55280 | Cell division protein FtsZ homolog 1 |
| HTR12 | SALK_204227C | At1g01370 | Histone H3-like centromeric protein HTR12 |
| MAP65-1 | SALK_006083C | At5g55230 | Microtubule-associated protein 65-1 |
| MAP65-3 | SALK_147277C | At5g51600 | Microtubule-associated protein 65-3 |
| RANGAP1 | SALK_123512C | At3g63130 | RAN GTPase-activating protein 1 |
| TAN | SALK_033437C | At3g05330 | TANGLED1 |
| TPXL4 | SALK_076598C | At5g07170 | TPX2-like Group A |
| TPXL8 | SALK_099761C | At5g62240 | TPX2-like Group A |
| TUBG1 | SALK_019182C | At3g61650 | Gamma-tubulin 1 |
