## Supplemental table 2 for "Identification of Gene Targets for the Sprouting Inhibitor CIPC"

**Table S2**: Non-cell cycle-related Arabidopsis mutants tested for altered root growth response to CIPC

| **Orthologue name (where known)** | **T-DNA line ID** | **Locus** | **Description** |
| --- | --- | --- | --- |
| ADO3 | SALK_059480C | At1g68050 | Flavin-binding kelch repeat F box 1 |
| AGO2 | SALK_201709C | At1g31280 | Protein argonaute 2 |
| AGO3 | SALK_204035C | At1g31290 | Protein argonaute 3 |
| ARR4 | SALK_060547C | AT1G10470 | Two-component response regulator ARR4 |
| AT1G03400 | SALK_201228C | At1g03400 | 2-oxoglutarate (2OG) and Fe(II)-dependent oxygenase |
| AT1G53050 | SALK_203826C | At1g53050 | Protein kinase |
| AT2G30650 | SALK_206166C | At2g30650 | Probable 3-hydroxyisobutyryl-CoA hydrolase 2 |
| At2g30820 | SALK_027207C | At2g30820 | Aspartyl/glutamyl-tRNA amidotransferase subunit |
| AT2G45170 | SALK_126394C | At2g45170 | Autophagy-related protein 8e |
| AT5G38940 | SALK_206432C | At5g38940 | RmlC-like cupins superfamily protein |
| AT5G39180 | SALK_080302C | At5g39180 | Germin-like protein subfamily 1 member 19 |
| AT5G43440 | SALK_092021C | At5g43440 | 1-aminocyclopropane-1-carboxylate oxidase homolog 9 |
| AT5G44290 | SALK_208688C | At5g44290 | Protein kinase |
| ATG8G | SALK_205553C | At3g60640 | Autophagy-related protein 8g |
| CHI-B | SALK_207226C | AT3G12500 | Basic endochitinase B |
| CUS2 | SALK_205690C | At5g33370 | Cutin synthase 2 |
| CYP76G1 | SALK_072380C | At3g52970 | Cytochrome P450, family 76, subfamily G, polypeptide 1 |
| FLS2 | SALK_023235C | At5g63580 | Putative inactive flavonol synthase 2 |
| FLS4 | SALK_002309C | At5g63595 | Probable flavonol synthase 4 |
| FLS5 | SALK_203500C | At5g63600 | Flavonol synthase 5 |
| FTSHI2 | SALK_054712C | At3g16290 | ATP-dependent zinc metalloprotease FTSHI 2 |
| GLP6 | SALK_205452C | AT5G39100 | Germin-like protein 6 |
| GLP9 | SALK_203089C | At4g14630 | Germin-like protein 9 |
| KNAT3 | SALK_136464C | At5g25220 | Knotted1-like homeobox gene 3 |
| KTI1 | SALK_131716C | At1g73260 | Kunitz trypsin inhibitor 1 |
| MTPC4 | SALK_071330C | AT1G51610 | Metal tolerance protein C4 |
| NSE4A | SALK_202958C | At1g51130 | Non-structural maintenance of chromosomes |
| OSM34 | SALK_206932C | At4g11650 | Osmotin 34 |
| PER36 | SALK_001604C | At3g50990 | Peroxidase 36 |
| PSRP3/1 | SALK_104063C | AT1G68590 | 30S ribosomal protein 3-1, chloroplastic |
| RAD51C | SALK_021960C | At2g45280 | RAS associated with diabetes protein 51C |
| RPA2B | SALK_022326C | At3g02920 | Replication protein A subunit B |
