## Supplemental figure 1 for "Identification of Gene Targets for the Sprouting Inhibitor CIPC"

### Slide 1
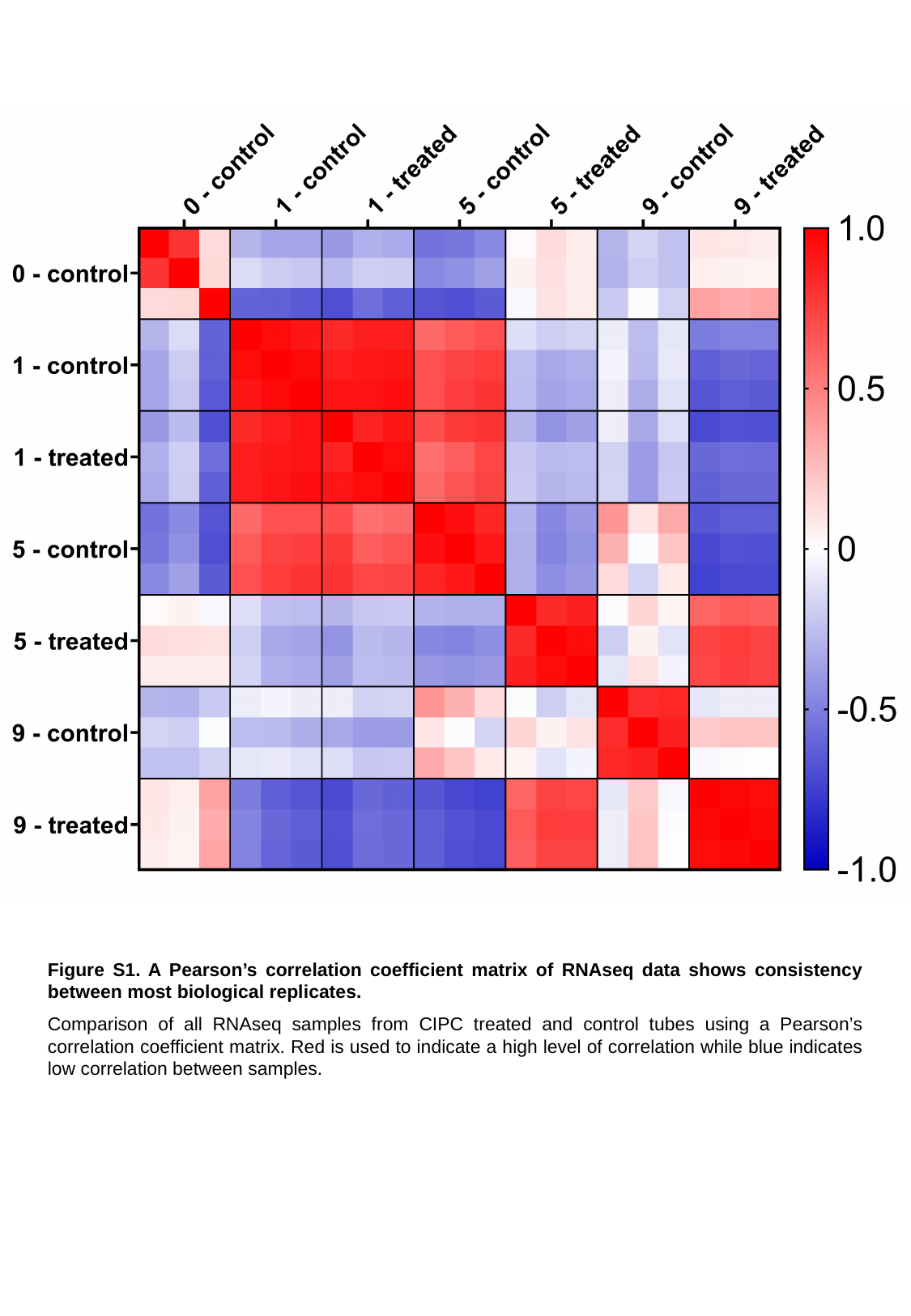

Figure S1. A Pearson’s correlation coefficient matrix of RNAseq data shows consistency between most biological replicates.
Comparison of all RNAseq samples from CIPC treated and control tubes using a Pearson’s correlation coefficient matrix. Red is used to indicate a high level of correlation while blue indicates low correlation between samples.
